# Evaluating the validity of the eye movement event detection model of Ganzin Sol glasses

**DOI:** 10.1101/2023.12.18.572270

**Authors:** Sung-En Chien, Kueian Lee, Chia-Yang Chang, Shaw-Chun Lu, Shao-Yi Chien

## Abstract

Eye tracking requires precise measurement of gaze data. Identifying fixations and saccades reliably is crucial for eye trackers, given these are fundamental eye movements. The present study evaluated the performance of Ganzin Sol glasses, a wearable eye tracker developed by Ganzin Technology, in detecting eye movement events. Participants performed the fixation and saccade invocation task (Komogortsev et al., 2010) and the gap paradigm (Saslow, 1967) using both Ganzin Sol and Tobii Pro 2 glasses separately at both short (50 cm) and long (300 cm) viewing distance. The fixation and saccade invocation task involved maintaining fixation on a regularly shifting visual target, enabling quantitative and qualitative analysis of participants’ oculomotor behaviors. In the gap paradigm, participants executed saccades toward a peripheral target when the fixation disappeared (gap condition) or remained visible (overlap condition). Typically, saccade latency in the gap condition is shorter (i.e., the gap effect). Results revealed aligned performances with Ganzin Sol and Tobii Pro 2 glasses in the fixation and saccade invocation task and the observation of the gap effect with both eye trackers. Therefore, the validity of Ganzin Sol glasses was at least comparable to Tobii Pro 2 glasses in the present study.

## Introduction

Eye tracking involves accurately measuring the point of gaze, and its applications span across various fields such as neuroscience, psychology, education, sports science, marketing, human-computer interaction (HCI), and product design.

For studies of oculomotor behavior, fixation and saccade are two primary eye movements (Hessels et al., 2018). Fixation is defined as the state when the eyes remain steady at a single target position over a period of time and is generally considered a reflection of attention (Holmqvist, 2011). Saccade refers to rapid changes between different target locations in the visual field (Leigh & Zee, 2015). These two measurements are commonly used in various research fields, including applied areas such as marketing and education. For example, marketing researchers might be interested in the distribution of consumer attention on a supermarket shelf (Bialkova et al., 2020). Educational researchers might also be interested in the degree of focused attention during a class (Jarodzka et al., 2021). Thus, it is important for an eye-tracker to reliably identify fixations and saccades from gaze data.

Two main types of methods have been proposed for classifying fixations and saccades: velocity- and position-based algorithms. The velocity threshold identification (I-VT) model (Duchowski, 2007; Salvucci & Goldberg, 2000) is one of the most representative velocity-based algorithms, which analyzes the velocity signal of the eye movement data. The I-VT model calculates the velocity for each gaze position sample, comparing it to a predefined threshold. If the calculated velocity is below the threshold, the corresponding gaze sample is classified as a fixational sample; otherwise, it is marked as a saccadic sample.

The dispersion threshold identification (I-DT) model is a commonly used position-based algorithm (Duchowski, 2007; Salvucci & Goldberg, 2000). The I-DT model defines a temporal window that shifts one gaze sample at a time. It evaluates the spatial dispersion created by the samples within this temporal window against a specified threshold. If the dispersion is below the threshold, the samples within the temporal window are categorized as fixational samples. Conversely, if the dispersion exceeds the threshold, the window advances by one sample, and the initial sample of the preceding window is designated as a saccadic sample.

### The goal of the present study

The Ganzin Sol glasses (Figure 1A) are a wearable eye-tracker developed by Ganzin Technology Inc (New Taipei City, Taiwan). The eye movement data analysis software, Caelus, can further identify fixations and saccades from the gaze samples using the I-VT model. Since the Sol glasses were only recently announced, it is necessary to evaluate the validity of the I-VT model implemented in the data analysis software to ensure that the quality of the recorded eye movement data matches the performance of other wearable eye trackers currently available on the market.

**Figure 1.**
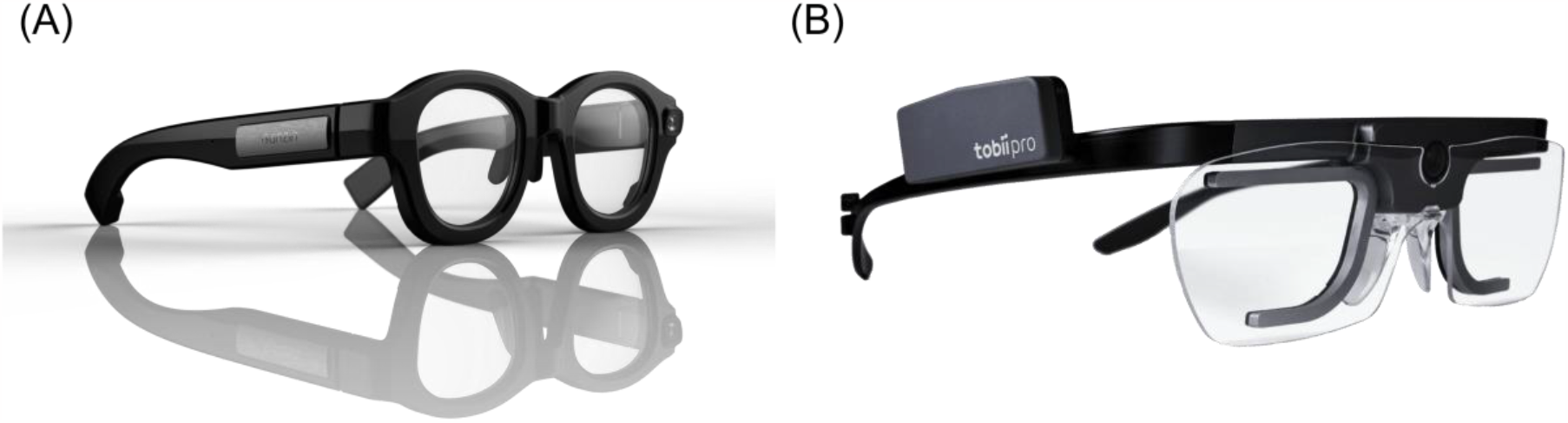
(A) Ganzin Sol glasses and (B) Tobii Pro 2 glasses.

To achieve this goal, the present study adopted a standardized task—the fixation and saccade invocation task proposed by Komogortsev et al. (2010)—and its derived metrics to evaluate the performance of the Ganzin Sol glasses in classifying fixations and saccades. These metrics include Fixation Quantitative Score (FQnS), Fixation Qualitative Score (FQlS), and Saccade Quantitative Score (SQnS). FQnS specifically assessed the classification algorithm’s capability to detect the amount of fixational behavior in response to given stimuli. FQlS examined the spatial proximity of fixational gaze samples to given stimuli. SQnS evaluated the algorithm’s effectiveness in detecting saccadic behaviors in response to given stimuli. Moreover, participants were instructed to perform the fixation and saccade invocation task not only with Ganzin Sol glasses but also with Tobii Pro 2 glasses (Figure 1B, Tobii AB, Stockholm, Sweden) — one of the most commonly used wearable eye trackers in studying human behaviors. We expected to observe comparable performances by participants in the fixation and saccade invocation task with Ganzin Sol glasses and those measured with Tobii Pro 2 glasses, assuming the performances of Ganzin Sol and Tobii Pro 2 glasses matched each other.

In addition, eye-tracking is widely used in studying human visual perception and attention. Such studies usually require accurate and stable data quality because the shifts of visual attention could happen in a short time window (e.g., within 1000 ms). Hence, we also adopted the classical gap paradigm (Saslow, 1967) and asked participants to perform a prosaccade task with both Ganzin Sol and Tobii Pro 2 glasses. In the gap paradigm, participants were instructed to perform saccades towards a peripheral target as soon as possible. A fixation was presented before the onset of the target. Before the target onset, the fixation might vanish (the gap condition) or remain visible (the overlap condition). Typically, saccade latency in the gap condition should be shorter than in the overlap condition, which is referred as the gap effect and could be explained by the facilitation of attentional disengagement induced by the disappearance of the fixation (Jin & Reeves, 2009). If the Ganzin Sol glasses are capable studying human visual perception, we expected that the gap effect could be observed with both Ganzin Sol and Tobii Pro 2 glasses.

## Method

### Participants

Sixteen participants (six females, mean age = 33.50, SD=7.13) were recruited from Ganzin Technology. Participants all had normal or corrected-to-normal visual acuity.

#### Apparatus

The stimuli were presented on a 24-inch LED screen (Dell U2414H, Dell Inc.) in the short-viewing-distance condition. In the long-viewing-distance condition, the stimuli were presented on a projector screen. Stimuli presentation was generated using C#. Participants’ eye movements were recorded by Ganzin Sol glasses (Ganzin Inc., New Taipei City, Taiwan) at 120-Hz sampling rate and Tobii Pro glasses 2 (Tobii AB, Stockholm, Sweden) at 100-Hz sampling rate.

### Stimuli, tasks, and procedures

Participants performed 1) the fixation and saccade invocation task (Komogortsev et al., 2010) and 2) the prosaccade task (Clark, 1999; Lee & Yeh, 2021; Pratt et al., 2006) in the present study.

In the fixation and saccade invocation task, a white disk served as the visual target against a gray background. The vertical position of the disk was consistently fixed in the central region of the screen. The disk had a diameter of 1°. At the beginning of the task, the disk appeared at the center of the screen for 1000 ms. Subsequently, the disk moved back and forth between left and right target positions. The left/right target positions were located 10° to the left or right of the screen’s center in the horizontal direction, repeated fourteen times. Participants were instructed to maintain fixation on the disk throughout the entire task. Consequently, there were a total of fifteen target positions, including the initial central fixation at the beginning of the task. (Figure 2A).

**Figure 2.**
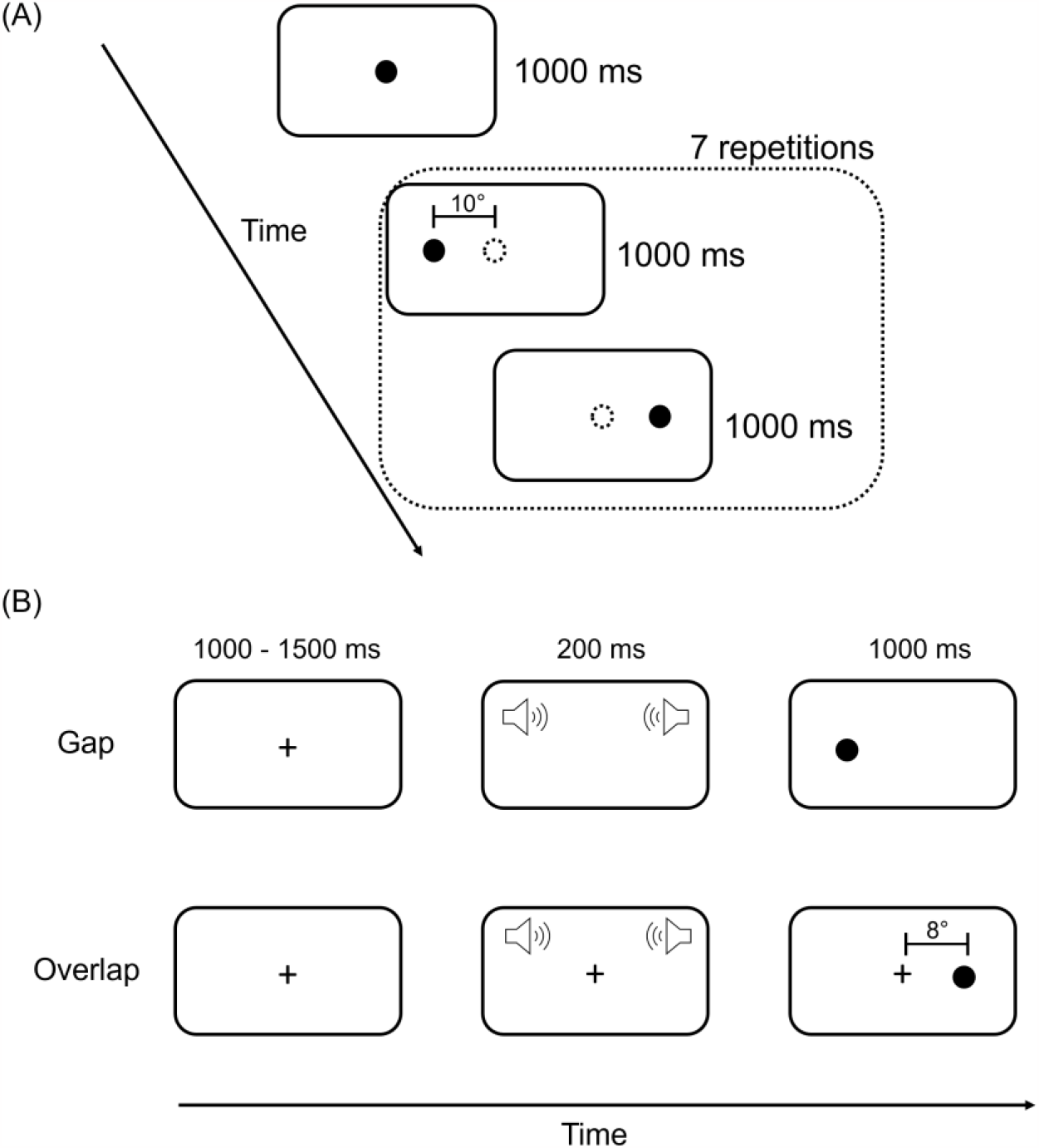
(A) Schematic presentation of the fixation and saccade invocation task. Participants were instructed to fixate at the target disk during the whole task. (B) Schematic presentation of the prosaccade task. Participants were asked to execute a saccade toward the target disk. The displays were not scaled.

In the prosaccade task, participants received instructions to fixate their gaze on a central fixation cross at the beginning of each trial. The dimensions of the fixation cross were both 1° in width and height. In each trial, the fixation cross would then appear at the center of the screen for a variable duration, ranging from 1,000 ms to 1,500 ms. There were two conditions in the prosaccade task: the overlap condition and the gap condition. In the overlap condition, the fixation cross remained on the screen for an additional 200 ms. Simultaneously, a white disk with a 1° diameter, referred to as the target, would appear either to the left or right of the fixation cross and remain visible for 1000 ms. The target’s positions were consistently located 8° away from the fixation cross (Saslow, 1967). In the gap condition, all parameters remained identical to those of the overlap condition, with the exception that the fixation cross disappeared earlier, and the screen remained blank for 200 ms (Figure 2B). In both conditions, a 1000 Hz tone was presented 200 ms before the target, serving as the cue for the target’s appearance. Participants were instructed to rapidly and accurately execute a saccade toward the target as soon as it became visible. There were 30 trials for both the overlap condition and the gap condition. Trials of different experimental conditions were presented in a randomized order.

For each participant, there were two different experimental conditions: the short-viewing-distance condition and the long-viewing-distance condition. In the short-viewing-distance condition, the viewing distance was 50 cm. Participants were situated in a quiet room, with their chin resting on a chin rest. In the long-viewing-distance condition, the viewing distance was 300 cm. The sizes of visual stimuli remained consistent in terms of visual angles in both the short- and long-viewing conditions. All participants first completed the short-viewing-distance condition, followed by the long-viewing-distance condition. Participants maintained standing position in the long-viewing-distance condition. In each condition, participants performed the fixation and saccade invocation task and the prosaccade task, using Ganzin Sol glasses and Tobii Pro glasses 2 separately. The sequence of the devices was counterbalanced across participants.

### Data analysis

#### Gaze samples

For both Ganzin Sol glasses and Tobii Pro 2 glasses, the descriptive statistics of the percentage of valid gaze samples in both the short-viewing distance condition and the long-viewing-distance condition are summarized in Table 1. The percentage was calculated by dividing the number of correctly identified gaze samples with usable gaze data by the total number of attempts. The valid sample ratios for all participants were consistently above 77%. The average valid sample ratios were consistently 90% or higher across all conditions.

**Table 1.** The descriptive statistics of the percentage of valid gaze samples.

| Device | Viewing distance | Mean | SD |
| --- | --- | --- | --- |
| Ganzin Sol | Short | 92% | 5% |
|  | Long | 95% | 3% |
| Tobii Pro 2 | Short | 90% | 5% |
|  | Long | 91% | 5% |

#### Eye movement events detection

For Tobii Pro 2 glasses, the Attention Filter in Tobii Pro Lab (Olsen, 2012) was used to detect fixations and saccades. In the case of Ganzin Sol glasses, fixations and saccades were identified using the IV-T model implemented by Ganzin Inc.

#### The fixation and saccade invocation task

We calculated the performance metrics (i.e., FQlS, FQnS, and SQnS) proposed by Komogortsev et al. (2010) to evaluate the validity of the Ganzin IV-T filter in the fixation and saccade invocation task and compared it with the Tobii Attention Filter.

FQnS was to compare the amount of detected fixational gaze samples to the total amount of gaze samples detected during the target presentation. Thus, the FQnS score represented the percentage of fixational gaze samples detected by the eye tracker. The ideal FQnS score should be 100%. However, since making a saccade toward the target position after the target onset took some times, the actual FQnS would be lower than 100%. Only when the recorded gaze sample was identified as a fixation with its position in spatial proximity to the target position, this gaze would be considered as a valid detected fixational gaze. The spatial proximity was set to 1/3 of the amplitude of the preceding stimulus saccade. The FQnS was calculated by normalizing the resulting detected fixational gaze samples by the total amount of gaze samples detected during the target presentation.

FQlS evaluated the spatial proximity between the detected eye fixation position and the presented target position, providing insights into the positional accuracy of the detected fixations. FQlS was calculated by summing the Euclidean distances in visual angle between matched pairs of target positions and the corresponding detected fixation positions throughout the task. This sum of Euclidean distances was then divided by the number of stimulus position points. Thus, the ideal FQlS should be closed to 0°.

SQnS represented the ratio between the amount of detected saccadic behaviors and the amount of saccadic behaviors encoded in the stimuli. This was computed by summing the absolute values of “detected saccade amplitude” and “stimulus saccade amplitude.” Subsequently, the total detected saccade amplitude was divided by the total stimuli saccade amplitude, and the resulting value was then normalized into a percentile. The ideal SQnS is 100%. The stimulus saccade was defined as the target’s movement to the next position. The detected saccades were those identified by the algorithm implemented in the eye-tracking system.

#### The prosaccade task

We calculated saccade latency in the prosaccade task. Saccade latency was defined as the time difference between the onset time of first saccade toward the target and the target onset time. Trials with saccade latencies shorter than 80 ms were also excluded to avoid saccade anticipation. In addition, trials with no saccade made toward the target were also excluded. Table 2 summarized the percentage of excluded trials in each condition.

**Table 2.** The summary of percentage the excluded trials in the prosaccade task.

| Device | Viewing distance | Excluded trilas (%) |
| --- | --- | --- |
| Ganzin Sol | Short | 13% |
|  | Long | 18% |
| Tobii Pro 2 | Short | 10% |
|  | Long | 13% |

## Results

### The fixation and saccade invocation task

#### FQnS

Figure 3A showed the results of the FQnS scores under different conditions. A two-way repeated-measure analysis of variance (ANOVA) on the FQnS scores was conducted on the factors of device type (Ganzin Sol glasses and Tobii Pro 2 glasses) and viewing distance (short-viewing-distance and long-viewing-distance). The main effect of device type was significant, *F*(1, 15) = 26.69, *p* < .001, η^2^_p_ = 0.64. The main effect of viewing distance was also significant, *F*(1, 15) = 7.54, *p* = .02, η^2^_p_ = 0.34. The interaction between the device type and the viewing distance was not significant, *F*(1, 15) = 1.37, *p* = .26, η^2^_p_ = 0.08 . Post-hoc analyses with Bonferroni correction showed that the FQnS scores measured with Ganzin Sol glasses were higher than the FQnS scores measured with Tobii Pro 2 glasses in both the short-viewing-condition (*t*(15) = 3.01, *p* = .03, Cohen’s *d* = 0.75) and the long-viewing condition (*t*(15) = 4.60, *p* < .001, Cohen’s *d* = 1.15).

**Figure 3.**
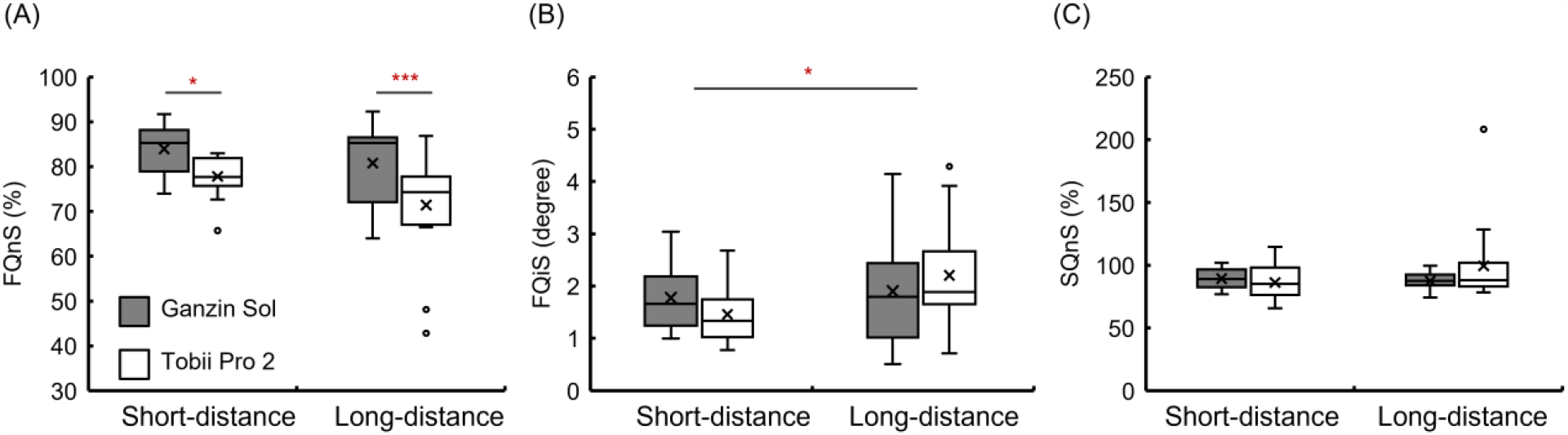
The results of the fixation and saccade invocation task (A) FQnS, (B) FQiS, and (C) SQnS. * and *** indicates *p*< .05 and *p*< .001, respectively. The error bar represents standard error.

#### FQlS

The results of the FQlS score were shown in Figure 3B. A two-way repeated-measure ANOVA was also conducted on the FQlS score on the same two factors as in the analysis of the FQnS score. The results showed that the main effect of device type was not significant, *F*(1, 15) = 3.79 × 10^-4^, *p* = .99, η^2^_p_ = 2.53 ×10^-5^. The main effect of viewing distance was significant, *F*(1, 15) = 6.17, *p* = .03, η^2^_p_ = 0.29, suggesting that the FQlS score in the short-viewing condition (mean = 1.62, SD = 0.58) was lower than in the long-viewing condition (mean = 2.05, SD = 0.95) regardless of the device types. There was no significant interaction between device type and viewing distance, *F*(1, 15) = 2.20, *p* = .16, η^2^_p_ = 0.13.

#### SQnS

Figure 3C showed the results of the SQnS scores. A two-way repeated-measure ANOVA was also conducted on the SQnS score on the same two factors as in the analysis of the FQnS score. The main effect of device type was not significant, *F*(1, 15) = 0.86, *p* = .37, η^2^_p_ = 0.05. The main effect of viewing distance was not significant, *F*(1, 15) = 1.75, *p* = .21, η^2^_p_ = 0.10. Although the interaction between device type and viewing distance was marginally significant, *F*(1, 15) = 3.29, *p* = .09, η^2^_p_ = 0.18, subsequent post hoc tests did not find any significant differences between conditions.

### The prosaccade task

Figure 4 showed the results of the prosaccade task. A three-way repeated-measure ANOVA on the saccade latency was conducted on the factors of device type, viewing distance, and fixation presentation (gap and overlap). The results showed that the main effect of device type was not significant, *F*(1, 15) = 0.004, *p* = .95, η^2^_p_ = 7.15 ×10^-5^. The main effect of view distance was not significant, *F*(1, 15) = 0.11, *p* = .75, η^2^_p_ = 0.01. The main effect of fixation presentation was significant, *F*(1, 15) = 61.40, *p* < .001, η^2^_p_ = 0.80. All two-way and the three-way interactions between factors were all insignificant. Hence, the results suggested that the saccade latency in the Gap condition (mean = 154.54, SD = 52.78) was less than in the overlap condition (mean = 193.20, SD = 48.32) regardless of device types and viewing distances.

**Figure 4.**
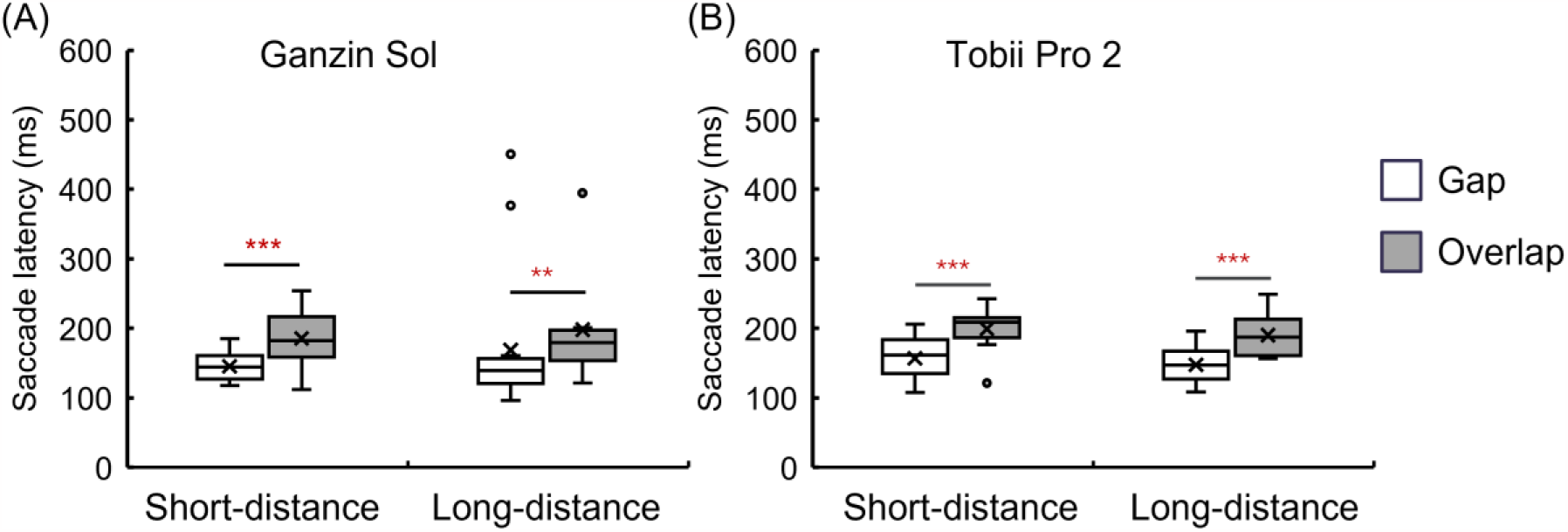
The results of the prosaccade task. ** and *** indicates *p*< .01 and *p*< .001, respectively. The error bar represents standard error.

Specially, saccade latencies measured with the Ganzin Sol glasses (Figure 4A) in the Gap condition were significantly less than the overlap condition in both the short-(*t*(15)= 5.96, *p* < .001, Cohen’s *d* = 0.78) and long-viewing-distance conditions (*t*(15)= 2.77, *p* = .002, Cohen’s *d* = 0.57). Similarly, with Tobii Pro 2 glasses (Figure 4B), saccade latencies in the Gap condition were also significantly less than the overlap condition in both the short-(*t*(15)= 6.27, *p* < .001, Cohen’s *d* = 0.82) and long-viewing-distance conditions (*t*(15)= 6.39, *p* < .001, Cohen’s *d* = 0.84).

## Discussion

### The fixation and saccade invocation task

The purpose of the present study was to examine the validity of the IV-T model of the Ganzin Sol glasses. To this goal, we asked the participants to perform the fixation and saccade invocation task and the prosaccade task with both short- and long-viewing distance with the Ganzin Sol glasses and the Tobii Pro 2 glasses, respectively.

In the fixation and saccade invocation task, the FQnS score measured with the Ganzin Sol glasses was higher the FQnS score measured with the Tobii Pro 2 glasses. suggesting that the Ganzin Sol glasses detected more fixational gaze samples when participants were asked to fixate their gazes on the target positions. In general, higher FQnS score reflected better performance of the eye tracker in detecting fixational behaviors. However, it should be noticed that there were outliers in FQnS score with Tobii Pro 2 glasses (Figure 3A). Since we didn’t measure FQnS scores with both eye trackers at the same time, and the ratios of valid gaze sample were generally high across participants and devices, it is still possible that the difference in FQnS scores between eye trackers was caused by actual differential performance when performing the fixation and saccade invocation task. For example, participants might blink more frequently when performing the task with Tobii Pro 2 glasses than with the Ganzin Sol glasses, resulting in less fixational samples detected.

The FQlS in the short-viewing-distance condition was generally lower than in the long-viewing-distance condition, suggesting that better positional accuracy of the detected fixations. That is, for both eye trackers, positional accuracy decreased with the increased viewing distance despite the distance between the target positions in visual angle remained the same in the long-viewing-condition as in the short-viewing condition.

The were no significant differences in SQnS between all conditions, suggesting that the amount of saccadic behaviors detected with Ganzin Sol glasses was comparable to the those detected with Tobii Pro 2 glasses regardless of viewing distances.

### The prosaccade task

The saccade latency in the Gap condition was less than the saccade latency in the Overlap condition across different device types and viewing distance, suggesting that we were able to observe the classical gap effect with both eye trackers. The gap paradigm requires stable temporal accuracy to reveal the subtle differences in reaction time of shifting gaze toward the target position induced by the absence/presence of the fixation in the Gap/Overlap condition. The successful replication of the gap paradigm indicated that the Ganzin Sol glasses is also capable of measuring the eye movement behaviors in behavioral experiments conducted in the laboratory settings.

## Conclusions

In brief, in the fixation and saccade invocation task, we observed more amount of detected fixational gaze samples with the Ganzin Sol glasses compared to the Tobii Pro 2 glasses. There was no difference in regard to the positional accuracy of the detected fixations and the amount of detected saccadic behaviors between the two different eye tackers. The classical gap effect could also be observed with both eye trackers. Thus, the validity of the Ganzin Sol glasses was at least comparable to the Tobii Pro 2 glasses in the present study.

## Notes

### Competing Interest Statement

All authors are employees of Ganzin Technology Inc.

